# Linked deterioration of early visual perception, function and structure in healthy human aging

**DOI:** 10.1101/2020.08.05.238014

**Authors:** Maria Fatima Silva, Ben M Harvey, Lília Jorge, Nádia Canário, Fátima Machado, Mário Soares, Miguel Castelo-Branco

**Affiliations:** Coimbra Institute for Clinical and Biomedical Research (iCBR), Faculty of Medicine, University of Coimbra, Coimbra, Portugal; Coimbra Institute for Biomedical Imaging and Translational Research (CIBIT), University of Coimbra, Coimbra, Portugal; Experimental Psychology, Helmholtz Institute, Utrecht University, Utrecht, The Netherlands; Institute of Nuclear Sciences Applied to Health (ICNAS), University of Coimbra, Coimbra, Portugal; CHUC, Ophthalmology Department, University of Coimbra, Coimbra, Portugal

**Author notes:** These authors contributed equally (Maria Fatima Silva, Ben M Harvey). Corresponding Author, Address: CIBIT-ICNAS, University of Coimbra, Azinhaga de Santa Comba, 3000-548, Coimbra, Portugal.

**Keywords:** Visual cortex, Aging, Retinal thickness, Visual acuity, fMRI, Population receptive field (pRF) modeling

## Abstract

Low-level visual perception deteriorates during healthy aging. We hypothesized that age-related retinal and cortical structure deteriorations affect perception through specific disruptions of neural function. We measured perceptual visual acuity in fifty healthy adults aged 20-80 years. We then measured these participants’ early visual field map (V1, V2 and V3) functional population receptive field (pRF) sizes and structural surface areas using fMRI, and their retinal structure using high-definition optical coherence tomography. With increasing age visual acuity decreased, pRF sizes increased, visual field maps surface areas decreased, and retinal thickness decreased. Among these measures, only functional pRF sizes predicted perceptual visual acuity. PRF sizes were in turn predicted by cortical structure only (surface areas), which were only predicted by retinal structure (thickness). We propose that age-related retinal structural deterioration disrupts cortical structure, thereby disrupting cortical functional neural interactions that normally sharpen visual position selectivity: the resulting functional disruption underlies age-related perceptual deterioration.

## 1. Introduction

Low-level visual perceptual abilities, like visual acuity, decline during healthy human aging. Aging is associated with structural changes in the retina including a gradual loss of retinal ganglion cells and their axons^1,2,3^. In the early visual cortex, primary visual cortex (V1) changes in both structure (decreasing surface area) and function (increasing receptive field sizes) between young and older adults^4,5^. We asked how these changes in the perception, structure and function of the early visual system in healthy human aging are linked in a large group of 50 healthy human subjects from 20 to 80 years of age. We hypothesized that age-related changes in retinal and cortical structure affect perception through specific changes in neural function.

The brain analyses visual space in a network of areas where the structural organization of the retina is repeatedly mapped onto the cortical surface^6,7,8^. This retinotopic structural organization of early visual areas V1, V2, and V3 has been well characterized in humans^9,10^. Recently developed functional magnetic resonance imaging (fMRI) methods also allow the functional response selectivity of neural populations to be characterized non-invasively in vivo. This approach, population receptive field (pRF) modeling^11^, relies on the gradual progression of single-neuron receptive field positions within retinotopic visual field maps, grouping together similarly-responding neurons. This is a major advance beyond localizing responsive areas and characterizing their structure: it allows comparisons of functional neural response properties between humans and even comparison of neural response selectivity to behavioral measures of perceptual abilities from the same subjects.

The visual position selectivity of neural populations (pRF size) in the early visual field maps has been well characterized in young adults. PRF sizes increase with visual eccentricity within a visual field map, and increase hierarchically between visual field maps. Smaller pRF sizes imply a finer neural representation of visual space. The last few years have seen an expansion of studies linking human visual perceptual abilities to functional pRF properties. PRF sizes are smaller where visual acuity is higher, near the foveal regions of the cortex^11,12^ and in the horizontal visual quadrants^13^, reflecting the spatial resolution of visual processing. The fine scale details of structural retinotopic organization within these visual field maps shows complementary changes. The cortical magnification factor (CMF, the cortical area responding to each degree of visual angle) increases in foveal regions^9,12^ and horizontal quadrants^13^, similarly implying a finer representation of visual space here. Recent studies in young adults have linked differences in pRF size and/or CMF to differences in perceptual performance across the visual field^13,14^ and between individual adults^15,16^.

It has been proposed that retinal ganglion cell loss may lead to receptive field enlargement^17,18^ in glaucoma models, through changes in cortical pooling mechanisms^19^ or a degradation of intracortical inhibition. We have recently used pRF properties to relate quality of vision to neural plasticity after cataract surgery in older adults^20,21^.

We therefore hypothesized that age-related loss of macular retinal ganglion cells may lead to decreased visual field map sizes, disrupting neural interactions and thereby increasing pRF sizes in the cortical central visual field representation. We further hypothesized that this increase in receptive field sizes ultimately underlies the common deterioration of visual acuity with age. To test this hypothesis, we combined perceptual measures of visual acuity with two neuroimaging methods: pRF modeling of fMRI data to quantify cortical visual field map structure and function, and high-definition optical coherence tomography (OCT) to quantify retinal structure. We tested the same 50 participants in all three data sets, spanning an age range from 20 to 80 years. This allowed us to quantify age-related changes in visual acuity (perception), early visual field map receptive field sizes (cortical functional) and surface areas (cortical structure), and retinal thickness (retinal structure). It also allowed us to examine between-participant associations in these measures to link the perceptual, functional and structural changes during healthy human aging throughout the early visual system.

## Results

### Visual acuity decreased with age

We quantified best corrected visual acuity (BCVA) using binocular ETDRS letter score (Figure 1A), where higher values indicate better vision. BCVA was significantly negatively correlated with age, with letter score decreasing by 1.25 points per decade (Figure 1B, statistics shown on figures).

**Figure 1:**
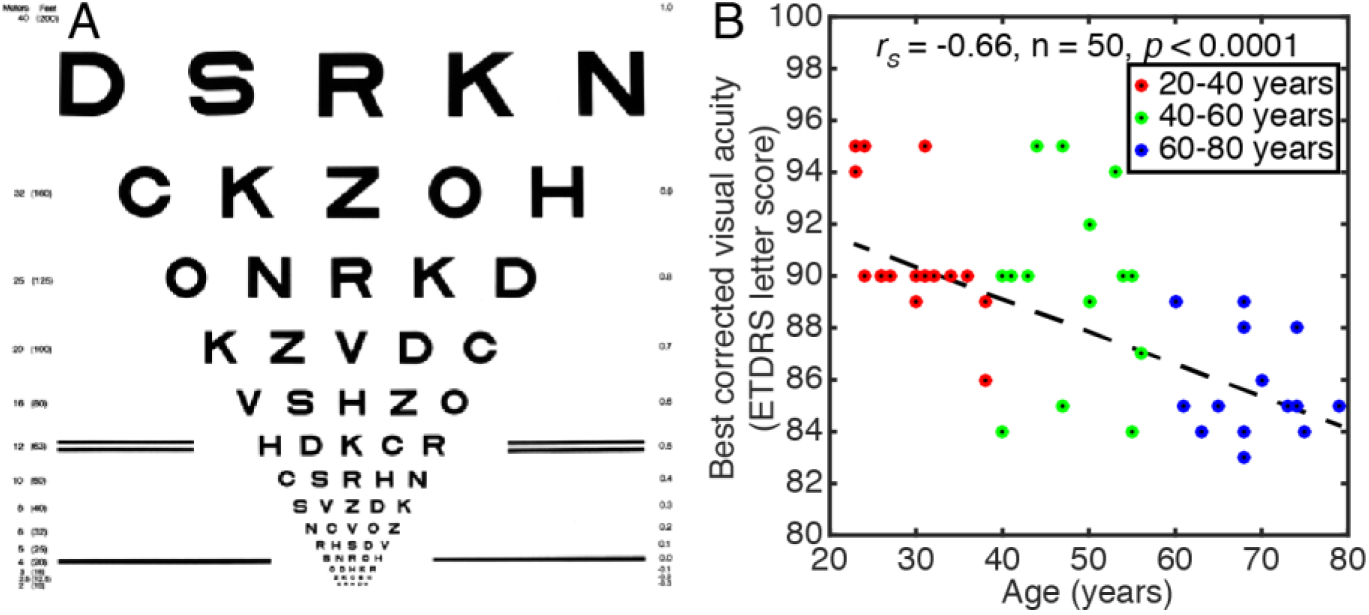
Visual acuity decreased with age. (A) Example test card for measuring best corrected visual acuity, from the National Eye Institute, National Institutes of Health. (B) Acuity decreased with age. Points are individual participants, dashed line is best linear fit.

### Visual field map pRF sizes and surface areas changed with age, and were correlated, but only pRF sizes predicted acuity

Maps of preferred visual field position polar angle and eccentricity across the early visual cortex were taken from pRF models for each participant. Figure 2A-B shows a representative hemisphere from young (20-40 years), middle-aged (40-60 years) and older (60-80 years) adults. All ages showed normal organization in these visual field maps. The visual field maps (V1, V2 and V3) were manually delineated for each participant.

**Figure 2:**
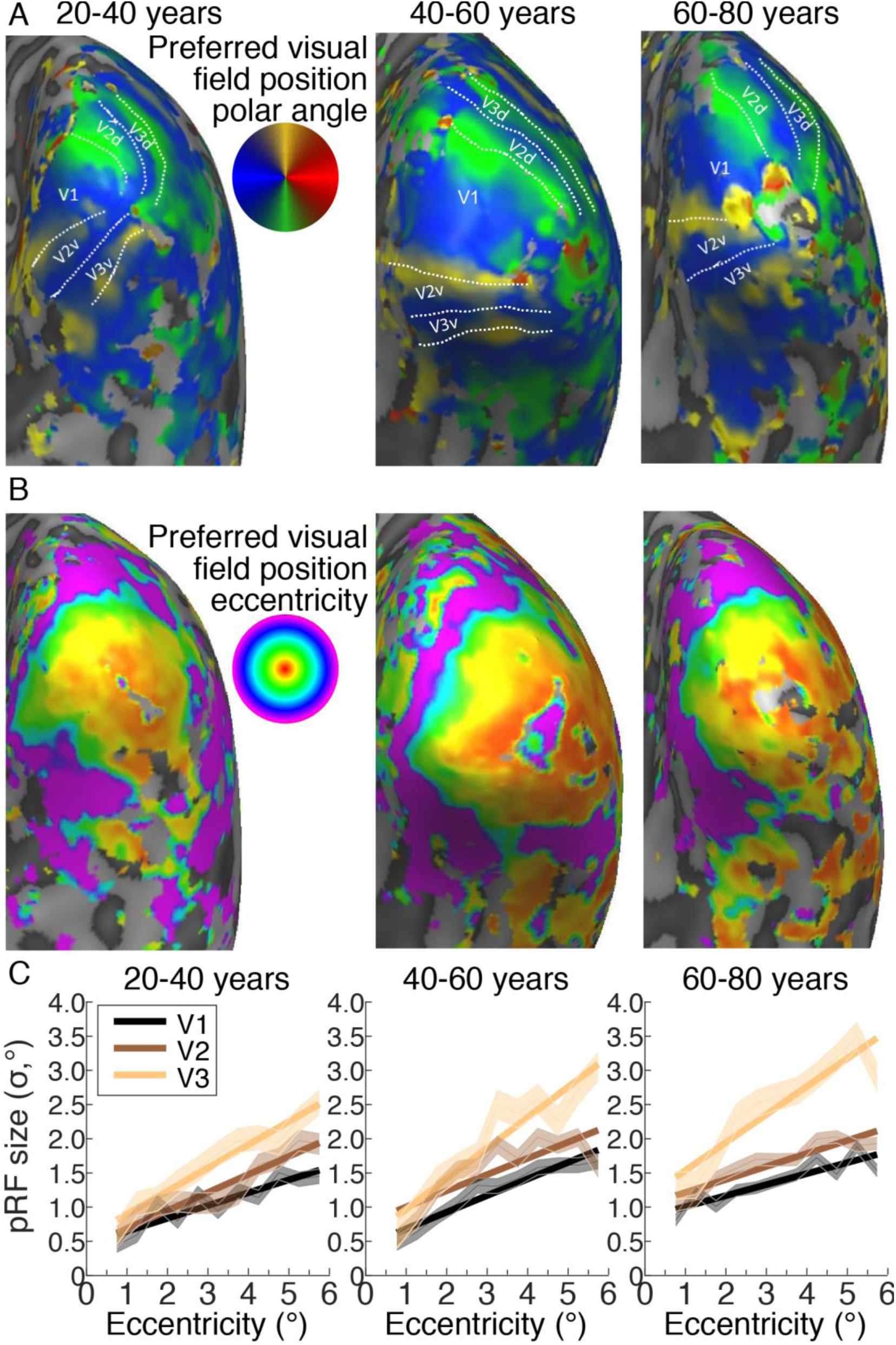
Visual field maps and pRF size changes with eccentricity across the early visual cortex. (A) Polar angle maps displayed on the inflated mesh (right hemisphere) for a representative example of each age group. The colors represent the recording sites for which the pRF model explains at least 30% of the variance. White dotted lines and labels show the position of the identified visual field maps. (B) Eccentricity maps displayed on the right hemisphere inflated mesh of representative examples of each age group. (C) PRF size changes with eccentricity in V1, V2 and V3 for a representative example participant from each age group. Shaded areas show the mean ± 1 SEM within each 0.5° eccentricity bin. Solid lines represent the best linear fit to bin means.

We then binned the pRF size of the recording sites in each visual field map into eleven eccentricity bins, each 0.5° wide, from 0.5° - 6°. Figure 2C shows changes in pRF size with eccentricity in V1, V2 and V3 representative example participants from each age group. To summarize these for each participant, we took a mean of these bins (global pRF size). We did not use the mean of individual recording site pRFs to avoid conflating pRF sizes with cortical magnification factors: cortical magnification decreases with eccentricity, so low eccentricities will contribute more to mean pRF sizes and differences in the distribution of cortical magnification factors will change the mean pRF size. Global pRF size avoids these issues. Global pRF sizes increased with age, showing strong correlations with age in V1, V2 and V3 (Figure 3A). PRF sizes increased by 0.11° in V1, 0.09° in V2 and 0.08° in V3 per decade. Global pRF sizes were strongly negatively correlated with visual acuity (Figure 3B), as previously shown within a narrower age range (Song et al., 2015).

**Figure 3:**
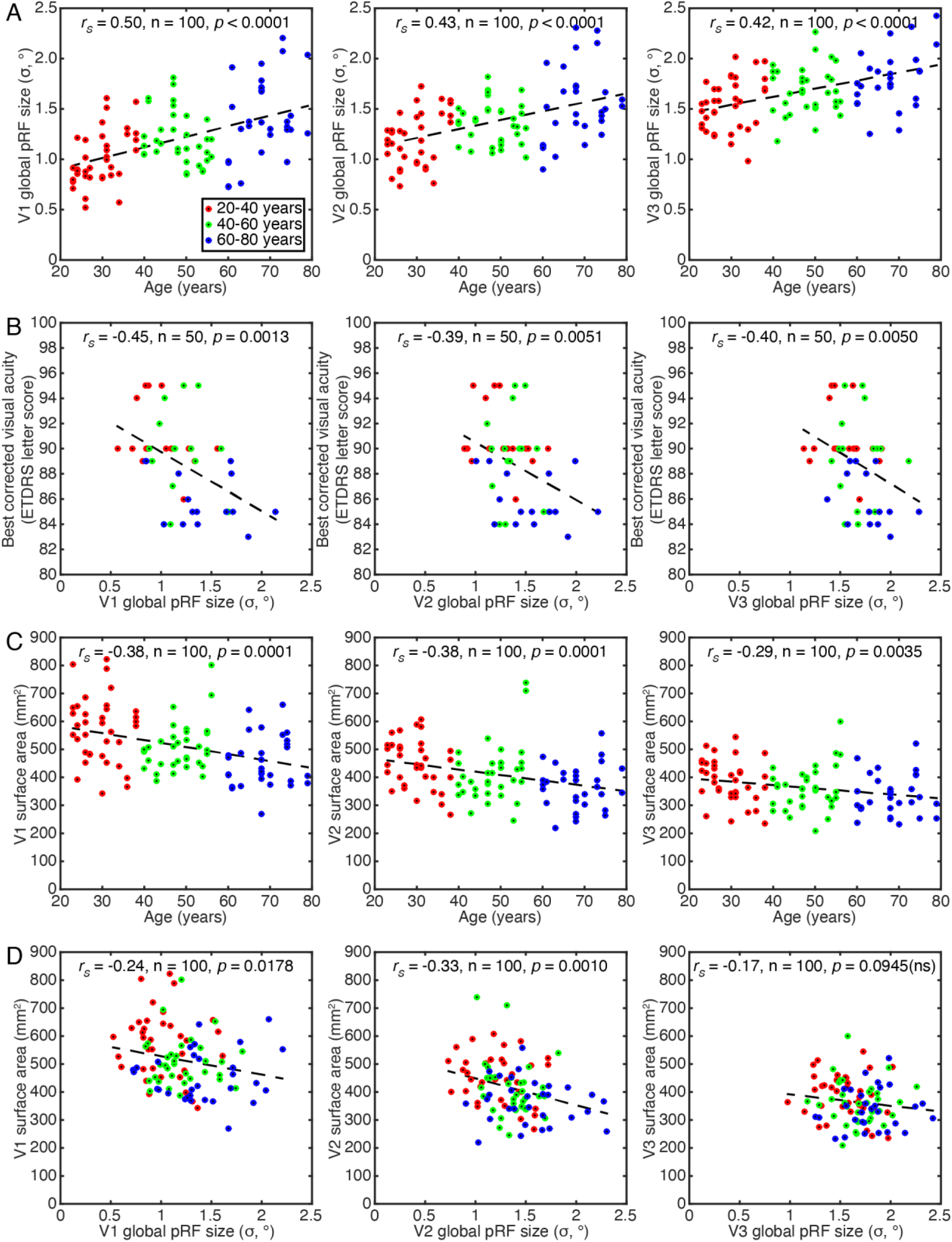
Age-related changes in global pRF size, surface area and their relationships. (A) Global pRF size increased with age. (B) BCVA decreased with increasing global pRF size. (C) Visual field map surface area decreased with age. (D) Visual field map surface area decreased as global pRF size increases in V1 and V2. Points are individual hemispheres (individual subjects in B), dashed line is the best linear fit.

Next, we asked whether global pRF sizes in different visual field maps were correlated across subjects. While not specifically relevant for questions of age and acuity, we can find no previous test of this relationship. We found strongly significantly correlations between pRF sizes in all visual field map pairs (V1 and V2: r = 0.797, *p* < 0.0001; V2 and V3: r = 0.708, *p* < 0.0001; V1 and V3: r = 0.583, p < 0.0001; all n = 100). To summarize each participant’s visual field map size, we measured the total surface area (in mm^2^) covered in each map from 0.5° - 6° eccentricity.

Visual field map surface areas decreased with age, showing strong negative correlations in V1, V2 and V3 (Figure 3C). Each hemisphere’s V1 shrank by 25 mm^2^ per decade, V2 by 19 mm^2^, and V3 by 12 mm^2^. So, age-related changes in both the visual field maps pRF sizes and surface area appear to decrease up the hierarchy. The surface areas of V1 and V2 were significantly negatively correlated with their pRF sizes (Figure 3D), while the correlation of V3’s surface area and pRF size only reached significance on a one-sided test (*p*=0.094 on a two-sided test). This correlation has been shown before in V1, within a narrower age range (Harvey and Dumoulin, 2011). However, no visual field map showed any significant relationship between its surface area and visual acuity (V1: *p*=0.084; V2: *p*=0.285; V3: *p*=0.790). Therefore, pRF sizes were more predictive of acuity than surface areas.

### Retinal thickness decreased with age and predicted visual field map surface areas

We measured global mean RT, GCIPLT, and RNFLT in all participants (Figure 4A). The measured values were consistent with normative data of Liu *et al*., (2011). All three measures were significantly negatively correlated with age (Figure 4B), with RT decreasing 2.2 µm, GCIPLT decreasing 1.5 µm and RNFLT decreasing 1.5 µm per decade. All three retinal measures were significantly correlated with the surface area of V1 (Figure 4C) and also V2 and V3 (Figure 5). However, no retinal measure was significantly correlated with pRF size (in any visual field map) or with visual acuity.

**Figure 4:**
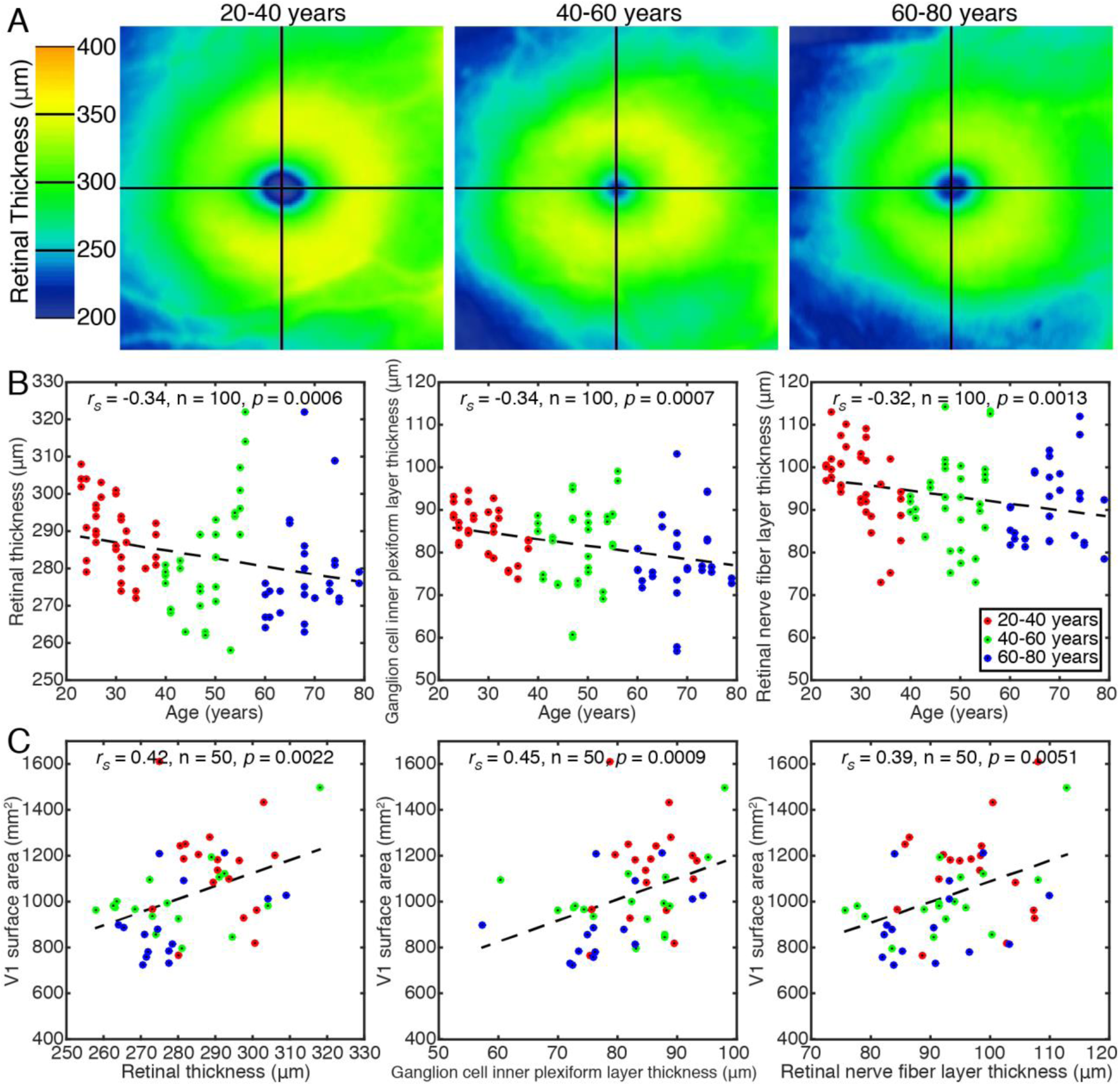
Age-related retinal thinning predicted decreases in V1 surface area. (A) Global retinal thickness (µm) maps for the central 20° of the right eye of representative participants from each age group. (B) The thickness of the total retina, ganglion cell inner plexiform layer and retinal nerve fiber layer decreased with age. (C) All retinal thickness measures predicted V1 surface area, but not pRF size or acuity. Points are individual eyes (in B) or participants (in C) dashed line is the best linear fit.

**Figure 5:**
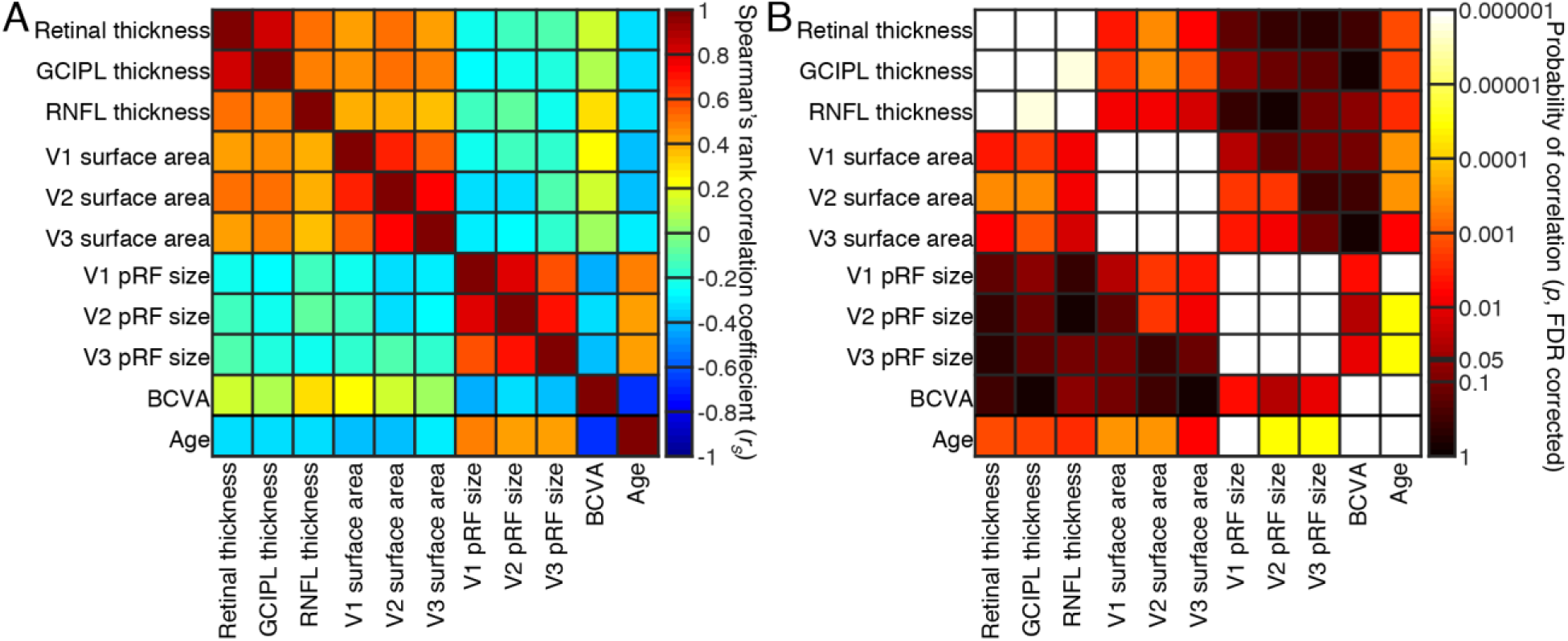
Correlations between all measures taken. (A) Pairwise Spearman’s correlation coefficients for all pairs of measures taken. (B) Probability of these correlations, after false discovery rate (FDR) correction for multiple comparisons.

### Integrated comparisons

The overall pattern of results so far suggests a series of linked processing steps: retinal thickness predicted visual field map surface areas, surface areas predicted pRF sizes, and pRF sizes predicted visual acuity. This is particularly evident in Figure 5B, where p-values of correlations become less significant (darker) with distance from the diagonal (i.e. with steps from the retina to perception), and no longer reached significance when measures are separated more than one of these steps.

However, we have tested this using separate correlations, and some of these measures are closely related and co-vary. Therefore, we also tested a set of general linear models of retinal thickness, V1 surface area, V1 global pRF size, and visual acuity, where each of these was used as an dependent variable, and the remaining three as independent predictors acting together. This demonstrated that V1 global pRF size and visual acuity significantly predicted each other (t=-3.45, df=46, p=0.0012), while other measures did not significantly predict either. Similarly, retinal thickness and V1 surface area significantly predicted each other (t=2.264, df=46, p=0.0054), while other measures did not significantly predict either. V1 surface area and pRF size did not predict each other here, apparently because the variance in these measures was better captured by retinal thickness and visual acuity respectively, to which they were more strongly correlated.

Nevertheless, significant correlations between V1 surface area and pRF size have been previously demonstrated (Harvey and Dumoulin, 2011), and were replicated here (Figure 3D).

## Discussion

In this study, we examined the neural basis of the common decline in visual acuity during healthy human aging by measuring visual acuity, retinal thickness, early visual field map surface areas, and their population receptive field (pRF) sizes in 50 adults from 20 to 80 years old. We characterized how these measures changed with age and how they co-varied. Retinal thickness, visual field map surface areas, and visual acuity all decreased with age, while pRF sizes increased. All these changes imply coarser visual processing. However, among these measures of neural structure and function, only functional pRF size significantly predicted visual acuity. PRF size was in turn predicted only by the visual field map’s surface area, which was in turn predicted by retinal thickness measures. These changes appear to be linked together such that, while all deteriorating with age, each step of the chain from the retina to the cortex to perception is only closely linked to the immediately preceding step. We therefore propose that structural deterioration of the retinal neural circuitry during healthy aging disrupts neural interactions that normally sharpen visual position selectivity, leading to an increase in cortical functional receptive field sizes through the early visual hierarchy, thereby decreasing visual acuity.

Our pRF size results are consistent with previously described increases in pRF size up the visual hierarchy, with recording sites’ preferred eccentricities, and in subjects with smaller V1 surface areas^12^. Previous results have also shown smaller pRF sizes in V1 and V2 predict lower perceptual position discrimination thresholds ^16^. These results all come from young and early middle-aged adults, 19-47 years old. Previous studies comparing 5 healthy young adults (24-36 years) and 4 healthy older adults (57-70 years) have shown larger pRF sizes and smaller V1 surface areas in the older group^4,5^. These two studies use the same data, from very small numbers of participants. This small sample size is concerning because pRF sizes and visual field map surface areas vary by a factor of two to three between healthy young adults^9,12^. We confirm this difference in our larger sample, and also show that pRF sizes are beginning to increase (and acuity decrease) in middle age. This suggests that age may be an important factor in individual differences in pRF size and acuity, and their covariation, even among the common sample of young and early middle-aged participants. However, the pRF size differences seen between young and middle-aged groups (an increase of ∼20%) are insufficient to explain the full range of individual differences (∼250%).

Regarding the retina, our OCT measures of retinal structure are also consistent with previous reports of gradual loss of retinal ganglion cells and their axons during healthy aging^1,2,3,22^. We propose that this retinal ganglion cell loss is likely to cause the decreases in visual field map surface areas, and (indirectly) the increases in pRF size, that we observe. For changes in visual field map surface areas, degradation of ascending retinal ganglion cell projections (in macular degeneration) causes a decrease in the gray matter volume in the affected cortical projection zone ^23^. Although V2 and V3 do not receive direct ascending projections, V2 is physically linked to V1 and V2 size is correlated with V1 size ^9^. Also, degradation of V1 seems likely to cause a similar atrophy of V2 and V3 through a reduction in feedforward activity. Interestingly, the shrinkage in V1 appears to occur faster (25 mm^2^ per decade) than in V2 (17 mm^2^ per decade), which is in turn faster than in V3 (17 mm^2^ per decade). This may suggest an effect in V1 being passed up the hierarchy.

Increases in early visual receptive field and pRF sizes are closely coupled to decreases in visual field map surface area or cortical magnification factor between individuals and within visual field maps^12,13,24^.However, retinal ganglion cell loss is also likely to indirectly affect receptive field sizes though cortical mechanisms: changes in cortical pooling^19^ or a degradation of intracortical inhibition, as seen in glaucoma models^17,18^.Previous results from senescent primates show a decrease (broadening) of orientation and motion direction selectivity in V1 and V2, coupled with increases in neural excitability^25,26,27^. These findings suggest an age-related degradation of intracortical inhibition resulting from the reliance of extrastriate receptive field properties on the upstream V1 receptive fields. As pRF sizes in extrastriate visual cortex (V2 and V3) are correlated with V1 pRF sizes, this increase in pRF size is likely to cascade through the visual hierarchy. Indeed, we see age-related increases in pRF size at least up to V3, with the change in pRF sizes again decreasing from V1 to V2 to V3 even as the pRF sizes themselves increase. Given that large pRF sizes in early visual field maps predict high visual position discrimination thresholds ^16^, such changes seem to underlie age-related reductions in visual acuity.

Together, our findings provide an integrated account of changes in perceptual visual acuity, retinal structure, and the structural organization and functional response selectivity of the early visual cortex during healthy aging. All of these measures are closely linked to age, but not all are closely linked to each other. We propose that deterioration of ascending retinal ganglion cells during healthy aging leads to a specific shrinkage of the cortical target of the ascending visual pathway, the primary visual cortex. This in turn disrupts cortical neural interactions that normally sharpen visual position selectivity, leading to an increase in cortical receptive field sizes that cascades through the early visual hierarchy. We propose that these changes in functional neural response selectivity are ultimately responsible for the age-related deterioration of visual perception: structural deterioration affects perception through specific effects on neural function. Therapies targeting the deterioration of the retinal ganglion cells may therefore prevent all these changes, so may be a promising approach to minimize the deterioration of visual perception during healthy aging.

## Methods

### Participants

Fifty healthy right-handed volunteers were recruited for this study and categorized by age into three groups: young adults (20-40 years, n = 18), middle-aged adults (40-60 years, n = 17), and older adults (60-80 years, n = 15). A full neuro-ophthalmological examination was performed, including best-corrected visual acuity (BCVA) measured as Early Treatment Diabetic Retinopathy Study (ETDRS) letter score using the ETDRS chart (higher scores correspond to better vision), intraocular pressure (IOP) measurement (Goldman applanation tonometer), slit-lamp biomicroscopy, fundus examination (Goldman lens). Retinal image acquisition was obtained with optical coherence tomography (spectral domain Cirrus HD-OCT 5000, Carl Zeiss Meditec, Dublin, CA, USA) and only participants without any abnormalities of the macula or the optic disc were included. All participants had normal or corrected to normal vision (visual acuity ≥ 8/10) and IOP ≤ 21 mmHg, and with no history of visual disease or clinical intervention. No subject showed any signs of Age-Related Macular Degeneration (even in an early stage), nor family history of glaucoma or other hereditary eye disease or diabetes. One subject was left-handed as determined by the Edinburgh inventor^28^ and excluded. Only participants without cognitive impairment were included in the study as assessed using the Montreal Cognitive Assessment-MoCA, a screening tool for cognitive deterioration^29^, scoring within normality according to their age and education. None of the subjects had an history of neurological or psychiatric disorders. The study was conducted in accordance with the tenets of the Declaration of Helsinki and was approved by the Ethics Committee of the University of Coimbra. Written informed consent for the study was obtained, after explanation of the nature and possible consequences of the study. Table 1 shows all demographic parameters of the study participants. There were no statistically significant differences between age groups in gender, education level and other demographic characteristic of participants.

**Table 1:** Participants’ demographic characteristics.

| Demographic parameters | Age groups |  |  |
| --- | --- | --- | --- |
|  | 20-40 years | 40-60 years | 60-80 years |
| Sample size (N subjects) | 18 | 17 | 15 |
| Mean age [*SEM*] (y) | 29.44 [1.15] | 48.24 [1.29] | 68.40 [1.51] |
| Age range (y) | 23-38 | 40 - 56 | 60 - 79 |
| Gender (male:female) | 9:9 | 11:6 | 6:9 |
| Mean weight [*SEM*] (Kg) | 66.11 [3.14] | 73.33 [4.12] | 67.04 [3.11] |
| Mean height [*SEM*] (Meters) | 1.68 [0.01] | 1.67 [0.02] | 1.67 [0.02] |
| Mean age of last education [*SEM*]: range (years) | 16.72 [0.61]:10-20y | 15.06 [1.01]: 6-20y | 13.79 [0.99]: 4-17y |

### fMRI Acquisition

Data acquisition was performed on a Siemens Magnetom Trio 3T scanner (Siemens, Erlangen, Germany), using a whole-brain approach, with a 12-channel head coil. Two high-resolution 3D anatomical MPRAGE (rapid gradient-echo) T1-weightened sequences were acquired, each with an isotropic resolution of 1 mm, repetition time (TR) of 2530 ms, echo time (TE) of 3.42 ms, field of view (FOV) of 256 × 256 mm. Each anatomical sequence comprised 176 slices, a flip angle of 7°, an inversion time of 1100 ms with a total time of 363 ms. The functional T2*-weighted 2D echo-planar MRI images were acquired with an isotropic resolution of 2 mm, TR of 2000 ms, TE of 30 ms, FOV of 256 ⨯ 256 mm. Each functional sequence comprised 29 interleaved slices and 186 volumes, the first 6 initial volumes for BOLD stabilization were discarded.

### Visual stimuli

Visual stimuli were displayed on a 32-inch LCD monitor (Inroom Viewing Device; NordicNeuroLab, Bergen, Norway) with 1920 ⨯1080 pixel resolution, positioned at the end of the scanner bore and viewed through a mirror attached to a head coil. The display was 70.0 × 39.5 cm and the viewing distance was 156.5 cm, so it subtended a 22.21° x 14.38° of visual angle.

The visual field mapping stimulus was created with PsychToolbox^30,31^ for Matlab (version R2014b; Mathworks, Natick, MA, USA). It consisted of bars stepping across the display perpendicular to bar orientation^11^, and revealing a white and black checkerboard pattern with 100% contrast moving parallel to the bar orientation (vertical, horizontal, and diagonal). The bar was 1.56° wide, 1/4^th^ of the stimulus radius, and the checks were each 0.78° square, so the spatial frequency of the checkerboard matched the bar width. Four bar orientations (0°, 45°, 90°, and 135°) and two different motion directions for each bar were presented giving a total of eight different bar motion directions, each of which crossed the display in 15 steps 0.625°, lasting 2 seconds each, 30 seconds. Four 30 second mean luminance (0% contrast) blocks were presented, one after each horizontal or vertical oriented bar crossing. Participant’s completed four visual field mapping runs (248 time frames each, around 6 min) within the same session.

A fixation dot at the center of the visual stimulus changed from red to green at random time intervals and participants were instructed to press a button on a response box every time they detected a color change, to ensure that attention and fixation were maintained. Color changes were every 3 s on average, with a minimum change interval of 1.8 s. We discarded any scan where detection performance dropped below 70% (2 scans of 1 subject). Mann-Whitney comparisons on the number of correct responses revealed no significant difference in performance between age groups (U = 28.000, *p* = 0.112).

### Anatomical and Functional Preprocessing

All fMRI data were processed and analyzed using BrainVoyager QX software (v2.8.4; Brain Innovation, Maastricht, the Netherlands). First, anatomical data underwent brain extraction and intensity inhomogeneity correction to reduce artifacts and inhomogeneity caused by the magnetic field ^32^. The two anatomic data sets were aligned to each other and in order to improve the signal-to-noise ratio, were averaged and re-oriented into AC-PC plane, followed by transformation to the Talairach reference system. The white matter was segmented by using an automatic segmentation routine^33^ and small manual adjustments were made, in order to create surface representations of each hemisphere (meshes). Preprocessing of the functional data included slice time correction, linear trend removal, temporal high-pass filtering (up to 2 cycles per scan), and 3D motion correction (rigid body) with spline interpolation. All volumes were corrected for head movement and motion artifacts between and within functional scans. Then, they were coregistered with each subject’s structural scan in Tailarach space and averaged across scans^34^.

### Population Receptive Field Estimation and Analysis

PRF models were estimated from the fMRI data and the time course of visual stimulus positions by using a model-driven approach^11^ and implemented in BrainVoyager QX. Shortly, this approach estimates a neural response model, for each voxel, that best explains each voxel’s fMRI response to the stimulus’s visual field positions. It models the preferred position (x and y) and size (standard deviation or σ) of a two-dimensional circular Gaussian function describing the area of the visual field to which the voxel responds. First of all, we generated a binary stimulus aperture containing the visual field positions covered by the stimulus bars in each TR. Next, for a large set of combinations of pRF positions and sizes, we calculated the proportion of the pRF function overlapping the stimulus aperture at every TR, proportional to the predicted neural response amplitude for this candidate pRF and this stimulus.

Next, each candidate neural response time course was convolved with a canonical BOLD hemodynamic response function to predict the BOLD response time course that the stimulus would yield for this set of pRF parameters. For each voxel, the pRF parameters (preferred position and size) were found that predict the BOLD response time course best correlated to the voxel’s response time course, with the variance explained by the model being equivalent to the R^2^ of this correlation.

The preferred position for each voxel was converted to preferred eccentricity and polar angle. These resulting parameter maps were projected onto inflated cortical meshes^35^, and the positions of V1, V2 and V3 were defined as regions of interest (ROIs) in relation to visual field representations^9,10,36^ (Figure 2) using BrainVoyager’s surface drawing tools. Voxels with pRF model variance explained below 0.3 were excluded from further analysis, as were voxels outside of the delineated ROI. We also excluded voxels with pRF eccentricities below 0.5°, since this part of the visual field is difficult to accurately map, and those beyond 6° eccentricity (near the edge of our stimulus area, where pRF properties are not estimated reliably. We attempted to delineate higher order areas, such as V4 and V3A, but in many participants these could not be identified reliably, because model fits were poor in those instances.

### Retinal imaging/thickness acquisition

Measurements of macular retinal thickness (RT), retinal nerve fiber layer thickness (RNFLT) and the ganglion cell-inner plexiform layer thickness (GCIPLT) were obtained with a Cirrus HD-OCT 5000 (Zeiss Meditec, Jena, Germany). All acquisitions were performed by the same trained operator. The volumetric data with 512 × 128 × 1024 voxels were acquired using the Macular Cube protocol centered on the macula. This protocol generates a cube of data through a 6 mm square grid around the fovea centralis by acquiring a series of 128 horizontal B-scans lines each composed of 512 A-scans, with an axial resolution of 5 mm. The standard output display of Cirrus HD-OCT includes a color topographical and a thickness map displaying measurements calculated for each of the nine macular areas corresponding to and defined by ETDRS grid. The global macular RT is defined as the average macular thickness from the inner limiting membrane to the top of the retinal pigment epithelium over the entire 6 × 6 scanned area. Global RT (mean of thicknesses in the nine sections of the ETDRS grid) were determined automatically and analyzed by the Cirrus HD-OCT’s internal algorithm. The thickness of two retinal layers was determined as well, such as RNFLT and GCIPLT. The RNFLT was acquired using the optic disc cube 200 × 200 protocol and analysis uses a 3.46 mm circle centered and around the optic disc. The ganglion cell analysis (GCA) algorithm measured the macular GCIPLT, within a 14.13 mm^2^ elliptical annulus area centered on the fovea. The global GCIPLT was measured in an elliptical annulus of the macular cube scan mode. Both eyes of each participant were separately scanned and compared, and no statistically significant differences between right and left eyes were found.

### Statistical Analysis

A MATLAB (version R2019b, Statistics and Machine Learning Toolbox) script was developed to extract the mean pRF size of each visual area for each participant. We then tested the effect of age on acuity (BCVA), visual field map pRF sizes, visual field map surface areas and retinal thickness measures using Spearman (non-parametric, rank) correlations. For consistency, we used non-parametric statistics throughout as some measures significantly deviated from normal distributions (in Shapiro-Wilk tests). Finally, we tested for relationships between these measures, again using Spearman correlations. Results with p<0.05 were considered statistically significant. We also tested a set of general linear models of retinal thickness, V1 surface area, V1 global pRF size, and visual acuity, where each of these was used as a dependent variable, and the remaining three as independent predictors acting together. Where two measurements are possible from each subject, either from the two eyes (retinal thickness) or the two hemispheres (pRF size, visual field map surface area), we treat these as two independent measurements. However, where testing for a correlation (or GLM) including a two-eye measure and a two-hemisphere measure, these pairs don’t match up. Here we use the mean retinal thickness from both eyes, the mean pRF size from both hemispheres, or the summed surface area from both hemispheres.

## Acknowledgements

This work was supported by the Portuguese Foundation for Science and Technology (#BPD/72029/2010 to MFS, BIGDATIMAGE, CENTRO-01-0145-FEDER-000016 financed by Centro 2020 FEDER, COMPETE; PAC – MEDPERSYST POCI-01-0145-FEDER-016428, #UID/NEU/0453904950/2013-2020 to MCB, #IF/01405/2014 to BH); and by the Netherlands Organization for Scientific Research (#452.17.012 to BH). The authors thank Bruno Direito, PhD, from the Coimbra Institute for Biomedical Imaging and Translational Research, for his collaboration.

## Author contributions

Conceptualization, MFS and MCB; Methodology, MFS and BH; Software, MFS and BH; Formal analysis, MFS and BH; Investigation, MFS, LJ, NC and MS; Data curation, MFS and FM; Writing - original draft, MFS and BH; Writing - review and editing, MFS, BH, LJ, NC MS and MCB; Visualization, MFS, BH and FM; Supervision, MCB; Project administration, MFS and MCB; Funding acquisition, BH and MCB.

## References

1. Pearson, P. M., Schmidt, L. A., Ly-Schroeder, E. & Swanson, W. H. Ganglion cell loss and age-related visual loss: A cortical pooling analysis. Optom. Vis. Sci. 83, 444–454 (2006). doi:10.1097/01.opx.0000218432.52508.10.

2. Harwerth, R. S., Wheat, J. L. & Rangaswamy, N. V. Age-related losses of retinal ganglion cells and axons. Investig. Ophthalmol. Vis. Sci. 49, 4437–4443 (2008). doi:10.1167/iovs.08-1753.

3. Harwerth, R. S. & Wheat, J. L. Modeling the effects of aging on retinal ganglion cell density and nerve fiber layer thickness. Graefe’s Arch. Clin. Exp. Ophthalmol. 246, 305–314 (2008). doi:10.1007/s00417-007-0691-5.

4. Brewer, A. A. & Barton, B. Effects of healthy aging on human primary visual cortex. Health (Irvine. Calif). 4, 695–702 (2012). doi:10.4236/health.2012.429109.

5. Brewer, A. A. & Barton, B. Visual cortex in aging and Alzheimer’s disease: Changes in visual field maps and population receptive fields. Front. Psychol. 5, 74 (2014). doi:10.3389/fpsyg.2014.00074.

6. Deyoe, E. A. et al. Mapping striate and extrastriate visual areas in human cerebral cortex. Proc. Natl. Acad. Sci. U. S. A. 93, 2382–2386 (1996). doi:10.1073/pnas.93.6.2382.

7. Wandell, B. A., Brewer, A. A. & Dougherty, R. F. Visual field map clusters in human cortex. Philos. Trans. R. Soc. B Biol. Sci. 360, 693–707 (2005). doi:10.1098/rstb.2005.1628.

8. Wandell, B. A. & Winawer, J. Imaging retinotopic maps in the human brain. Vision Research 7, 718–737 (2011). doi:10.1016/j.visres.2010.08.004.

9. Dougherty, R. F. et al. Visual field representations and locations of visual areas v1/2/3 in human visual cortex. J. Vis. 3(10), 1 (2003). doi:10.1167/3.10.1.

10. Wandell, B. A., Dumoulin, S. O. & Brewer, A. A. Visual field maps in human cortex. Neuron 56, 366–383 (2007). doi:10.1016/j.neuron.2007.10.012.

11. Dumoulin, S. O. & Wandell, B. A. Population receptive field estimates in human visual cortex. Neuroimage 39, 647–660 (2008). doi:10.1016/j.neuroimage.2007.09.034.

12. Harvey, B. M. & Dumoulin, S. O. The relationship between cortical magnification factor and population receptive field size in human visual cortex: Constancies in cortical architecture. J. Neurosci. 31, 13604–13612 (2011). doi:10.1523/JNEUROSCI.2572-11.2011.

13. Silva, M. F. et al. Radial asymmetries in population receptive field size and cortical magnification factor in early visual cortex. Neuroimage 167, 41–52 (2018). doi:10.1016/j.neuroimage.2017.11.021.

14. Duncan, R. O. & Boynton, G. M. Cortical magnification within human primary visual cortex correlates with acuity thresholds. Neuron 38, 659–671 (2003). doi:10.1016/S0896-6273(03)00265-4.

15. Schwarzkopf, D. S., Song, C. & Rees, G. The surface area of human V1 predicts the subjective experience of object size. Nat. Neurosci. 14, 28–30 (2011). doi:10.1038/nn.2706.

16. Song, C., Schwarzkopf, D. S., Kanai, R. & Rees, G. Neural population tuning links visual cortical anatomy to human visual perception. Neuron 85, 641–656 (2015). doi:10.1016/j.neuron.2014.12.041.

17. King, W. M. et al. Expansion of visual receptive fields in experimental glaucoma. Vis. Neurosci. 23, 137–142 (2006). doi:10.1017/S0952523806231122.

18. Sharma, S. C. Changes of central visual receptive fields in experimental glaucoma. Progress in Brain Research 173, 479–491 (2008). doi:10.1016/S0079-6123(08)01133-3.

19. Redmond, T., Garway-Heath, D. F., Zlatkova, M. B. & Anderson, R. S. Sensitivity loss in early glaucoma can be mapped to an enlargement of the area of complete spatial summation. Investig. Ophthalmol. Vis. Sci. 51, 6540–6548 (2010). doi:10.1167/iovs.10-5718.

20. Miranda, Â. S. C. et al. Optical properties influence visual cortical functional resolution after cataract surgery and both dissociate from subjectively perceived quality of vision. Investig. Ophthalmol. Vis. Sci. 59, 986–994 (2018). doi:10.1167/iovs.17-22321.

21. Rosa, A. M. et al. Functional Magnetic Resonance Imaging to Assess the Neurobehavioral Impact of Dysphotopsia with Multifocal Intraocular Lenses. in Ophthalmology 124, 1280–1289 (2017). doi:10.1016/j.ophtha.2017.03.033.

22. Jorge, L., Canário, N., Quental, H., Bernardes, R. & Castelo-Branco, M. Is the Retina a Mirror of the Aging Brain? Aging of Neural Retina Layers and Primary Visual Cortex Across the Lifespan. Front. Aging Neurosci. 11, 360 (2020). doi:10.3389/fnagi.2019.00360.

23. Hanson, R. L. W. et al. Following the status of visual cortex over time in patients with macular degeneration reveals atrophy of visually deprived brain regions. Investig. Ophthalmol. Vis. Sci. 60, 5045–5051 (2019). doi:10.1167/iovs.18-25823.

24. Hubel, D. H. & Wiesel, T. N. Uniformity of monkey striate cortex: A parallel relationship between field size, scatter, and magnification factor. J. Comp. Neurol. 158, 295–305. (1974) doi:10.1002/cne.901580305.

25. Leventhal, A. G., Wang, Y., Pu, M., Zhou, Y. & Ma, Y. GABA and its agonists improved visual cortical function in senescent monkeys. Science 300, 812–815 (2003).doi:10.1126/science.1082874.

26. Yu, S., Wang, Y., Li, X., Zhou, Y. & Leventhal, A. G. Functional degradation of extrastriate visual cortex in senescent rhesus monkeys. Neuroscience 140, 1023–1029 (2006). doi:10.1016/j.neuroscience.2006.01.015.

27. Schmolesky, M. T., Wang, Y., Pu, M. & Leventhal, A. G. Degradation of stimulus selectivity of visual cortical cells in senescent rhesus monkeys. Nat. Neurosci. 3, 384–390 (2000). doi:10.1038/73957.

28. Oldfield, R. C. The assessment and analysis of handedness: the Edinburgh Inventory. Neuropsychologia vol. 9 97–113 (1971).

29. Freitas, S., Simões, M. R., Alves, L. & Santana, I. Montreal Cognitive Assessment (MoCA): Normative study for the Portuguese population. J. Clin. Exp. Neuropsychol. 33, 989–996 (2011).

30. Brainard, D. H. The Psychophysics Toolbox. Spat. Vis. 10, 433–436 (1997).doi:10.1163/156856897X00357.

31. Pelli, D. G. The VideoToolbox software for visual psychophysics: Transforming numbers into movies. Spat. Vis. 10, 437–442 (1997). doi:10.1163/156856897X00366.

32. Dale, A. M., Fischl, B. & Sereno, M. I. Cortical surface-based analysis: I. Segmentation and surface reconstruction. Neuroimage 9, 179–194 (1999). doi:10.1006/nimg.1998.0395.

33. Kriegeskorte, N. & Goebel, R. An efficient algorithm for topologically correct segmentation of the cortical sheet in anatomical MR volumes. Neuroimage 14, 329–346 (2001). doi:10.1006/nimg.2001.0831.

34. Nestares, O. & Heeger, D. J. Robust multiresolution alignment of MRI brain volumes. Magn. Reson. Med. 43, 705–715 (2000). doi:10.1002/(SICI)1522-2594(200005)43:5<705::AID-MRM13>3.0.CO;2-R.

35. Wandell, B. A., Chial, S. & Backus, B. T. Visualization and measurement of the cortical surface. J. Cogn. Neurosci. 12, 739–752 (2000). doi:10.1162/089892900562561.

36. Sereno, M. I. et al. Borders of multiple visual areas in humans revealed by functional magnetic resonance imaging. Science 268, 889–893 (1995). doi:10.1126/science.7754376.

